# The lipid transfer protein Saposin B does not directly bind CD1d for lipid antigen loading

**DOI:** 10.1101/698654

**Authors:** Maria Shamin, Tomasz H. Benedyk, Stephen C. Graham, Janet E. Deane

## Abstract

Lipid antigens are presented on the surface of cells by the CD1 family of glycoproteins, which have structural and functional similarity to MHC class I molecules. The hydrophobic lipid antigens are embedded in membranes and inaccessible to the lumenal lipid-binding domain of CD1 molecules. Therefore, CD1 molecules require lipid transfer proteins for lipid loading and editing. CD1d is loaded with lipids in late endocytic compartments, and lipid transfer proteins of the saposin family have been shown to play a crucial role in this process. However, the mechanism by which saposins facilitate lipid binding to CD1 molecules is not known and is thought to involve transient interactions between protein components to ensure CD1-lipid complexes can be efficiently trafficked to the plasma membrane for antigen presentation. Of the four saposin proteins, the importance of Saposin B (SapB) for loading of CD1d is the most well-characterised. However, a direct interaction between CD1d and SapB has yet to be described. In order to determine how SapB might load lipids onto CD1d, we used purified, recombinant CD1d and SapB and carried out a series of highly sensitive binding assays to monitor direct interactions. Using equilibrium binding analysis, chemical cross-linking and co-crystallisation experiments, under a range of different conditions, we could not demonstrate a direct interaction. This work establishes comprehensively that the role of SapB in lipid loading does not involve direct binding to CD1d. We discuss the implication of this for our understanding of lipid loading of CD1d and propose several factors that may influence this process.

## Introduction

The presentation of antigenic molecules is fundamental to host defence and immune regulation. The MHC class I and II molecules present peptide antigens to T cells and the molecular mechanisms of peptide loading and editing are relatively well understood^1–3^. However, the mechanisms by which lipid antigens are loaded onto the CD1 family of antigen-presenting molecules remains poorly understood. CD1 molecules have structural similarity to MHC class-I, including association with β-2-microglobulin (β_2_m), but unlike MHC molecules, CD1 molecules are non-polymorphic and the antigen-binding grooves of CD1 molecules are extremely hydrophobic, allowing the binding of most classes of lipids including phospholipids, glycerolipids, lysolipids and glycolipids possessing a huge variety of glycosylated head groups^4^.

Of the five CD1 family members, antigen presentation by CD1d is best characterised. CD1d undergoes complex cellular trafficking in a manner similar to that of MHC class II. Following synthesis in the endoplasmic reticulum, CD1d traffics via the Golgi apparatus to the plasma membrane, and is then internalised and trafficked through early and late endocytic compartments before being returned to the plasma membrane^5^, where CD1d-lipid complexes are recognized by the T cell receptor (TCR) of natural killer T (NKT) cells. Invariant NKT (iNKT) cells, which bear an invariant TCR and are characteristically potently activated by recognition of CD1d loaded with the synthetic glycolipid α-galactosylceramide (α-GalCer), are innate-like T cells involved in the orchestration of immune responses^6^.

Lipid antigens are embedded in cellular membranes^7^ and thus are not directly accessible to the lumenal, lipid-binding groove of CD1 molecules. The loading of lipid antigen onto CD1 molecules therefore requires the action of lipid-transfer proteins (LTPs) for efficient antigen presentation. The loading of lipids onto CD1d can occur in the ER, at the cell surface and in endocytic compartments. Relatively little is known about how lipid specificity is determined, and studies using secreted or recycling, surface-cleavable human CD1d found differences in the lipid repertoire bound to CD1d, reflecting the lipids present in the compartments through which it had trafficked^8–10^. Different LTPs are present in the different cellular locations where lipid antigen loading occurs and are likely to determine the lipid repertoire bound to CD1 molecules^11^. Interestingly, the late endocytic/lysosomal compartment is the site of both glycolipid catabolism and lipid loading of CD1d and the resident LTPs function in both these processes. Importantly, the antigenicity of glycolipids can be regulated by processing of their glycan headgoups prior to presentation to T cells, highlighting the complex interplay between these two pathways^12,13^.

The LTPs present in the lysosomal compartments include ganglioside monosialic 2 activator (GM2A), Niemann-Pick type C 2 (NPC2) and four saposin (Sap) proteins. SapA, SapB, SapC and SapD are the products of the proteolytic cleavage of a prosaposin (PSAP) precursor and are each small, non-enzymatic proteins possessing three disulphide bonds that stabilise an extremely heat-resistant helical structure. The saposins can exist in a “closed” monomeric, globular conformation or, at low pH, adopt a more “open” conformation, forming higher-order oligomers enclosing a hydrophobic cavity into which lipid acyl chains can be buried^14–19^. Each saposin functions in conjunction with specific hydrolases to facilitate the degradation of different glycosphingolipids and the loss of saposin function phenotypically resembles the loss of the associated hydrolases^20–25^. There are two proposed mechanisms for how saposins assist in lipid presentation to hydrolases: the “solubiliser” model, whereby saposins encapsulate lipids within a hydrophobic oligomeric complex for presentation to soluble enzymes, and the “liftase” model, where saposins bind directly to membranes destabilising them allowing membrane-associated hydrolases access to lipid substrates. Structural and cell-based evidence exists to support both these models^19,26–29^. These two models are also relevant for the proposed mechanisms by which saposins load lipids onto CD1 molecules^30^. SapB is thought to behave as a “solubiliser”, forming protein-lipid complexes that enhance lipid loading of CD1 molecules, while SapC associates with membranes and may facilitate lipid loading at the membrane surface.

The evidence for saposin-mediated lipid loading of CD1 molecules comes from animal knockout studies, cell-based assays and *in vitro* assays. Knockout of prosaposin (PSAP) in mice results in the dramatic loss of presentation of endogenous iNKT cell ligands by CD1d^31^. The presentation of exogenous lipid antigens, such as α-GalCer, is impaired in PSAP knockout or knockdown cells^31–34^. *In vitro* assays have identified that all the saposins, as well as the other lysosomal LTPs, GM2A and NPC2, are capable of transferring lipids onto CD1 molecules^31,35^. However, some LTPs appear to be more efficient than others at loading specific lipid antigens onto different CD1 molecules^30,33,36^. Specificity within the saposin family in this process has been explored via *in vitro* assays identifying SapB as the dominant saposin for the loading of lipid antigens onto CD1d. Specifically, incubation of CD1d with SapB and α-GalCer has been shown to enhance the stimulation of iNKT cells^33,34^ and SapB can mediate lipid binding to CD1d in T cell independent assays^33^ suggesting a direct role for this specific saposin in lipid loading of CD1d. The ability of SapB to facilitate and enhance lipid exchange on CD1d has also been monitored *in vitro* via iNKT TCR binding to CD1d following incubation with lipids alone or lipids plus SapB^34^. These *in vitro* assays implicate SapB as a “lipid editor” that facilitates the loading and unloading of lipid antigens onto CD1d.

However, there have been several references in the literature to an inability to measure a direct interaction between saposins and human CD1d^11,33,34^. Unfortunately, these reports do not show any data or detail the approaches used to monitor binding and the implication is that these interactions exist but are weak and very transitory. Therefore we have used a number of robust and sensitive biochemical and structural techniques to monitor direct interactions between SapB and CD1d. Using multiple approaches in a range of conditions we do not observe any evidence of a direct interaction between CD1d and SapB.

## Methods

### Cloning and cell line production

Codon-optimised cDNA was synthesized (GeneArt) encoding human β_2_m (residues 21-119, Uniprot P61769) and the extracellular domain of human CD1d (residues 20-301, Uniprot P15813) with a C-terminal hexahistiding tag (CD1d-H_6_). A modified version of the piggyBac target protein plasmid PB-T-PAF^37^ was constructed retaining the N-terminal secretion signal but with the Protein A fusion removed. CD1d-H_6_ and β_2_m were individually cloned into this modified vector, using *Spe*I and *Asc*I restriction-endonuclease sites, to produce PB-T-CD1d-H_6_ and untagged PB-T-β_2_m. For production of an inducible, stable co-expression cell line, HEK293-F cells were quadruple-transfected with PB-T-CD1d-H_6_, PB-T-β_2_m, PB-RN and PBase using a DNA mass ratio of 5:5:1:1 and transfected cells were selected with geneticin using the established protocol for the piggyBac expression system^37^.

Codon-optimised cDNA was synthesized (GeneArt) encoding the segment of human prosaposin corresponding to SapB (residues 195-274, Uniprot P07602) and subcloned into the bacterial expression vector pET-15b using *Nco*I and *Xho*I restriction endonuclease sites to produce untagged protein. SapB was also subcloned into the mammalian expression vector pHLsec^38^ using the restriction enzymes *Age*I and *Kpn*I to encode a C-terminally H_6_-tagged protein.

### Protein expression and purification

Protein expression of CD1d-β_2_m complex in the HEK293F co-expression cell line was induced with 2 µg/mL doxycycline and expression media was collected over a 2 month period. CD1d-H_6_ was purified from conditioned media by nickel affinity chromatography in phosphate-buffered saline (PBS) pH 7.4 and eluted using PBS containing 300 mM Imidazole. CD1d-β_2_m was further purified by size exclusion chromatography on a Superdex 200 column (GE Healthcare) equilibrated in cross-linking buffer (20 mM HEPES pH 7.0, 150 mM NaCl) for interaction assays or distinct buffers (described below) for crystallisation experiments. CD1d-β_2_m was concentrated to 2 mg/mL and stored at 4°C for up to one month.

To produce deglycosylated CD1d-β_2_m, cells were treated with 5 μM kifunensine at the same time as protein expression was induced. CD1d-β_2_m was purified by nickel affinity purification and eluted with 100 mM citrate pH 4. CD1d-β_2_m protein was buffer exchanged into 100 mM citrate pH 5.5, using repeated rounds of dilution and concentration using a centrifugal concentrator, and concentrated to 2 mg/mL and incubated with 3125 Units of Endoglycosidase H (Endo H)(NEB) per milligram of protein for 3 hours at room temperature. Deglycosylated CD1d-β_2_m was purified by size exclusion chromatography (as described above).

Untagged SapB was expressed in *Escherichia coli* Origami (DE3) cells and purified as described previously^39^. Briefly, cleared lysate was heat-treated and precipitated proteins were cleared by centrifugation and supernatant containing SapB was dialysed overnight in the presence of 20µg/mL DNAse against anion exchange buffer (50 mM Tris pH 7.4, 25 mM NaCl). SapB was further purified by anion exchange chromatography (HiTrap QSepharose column) followed by size-exclusion chromatography (HiLoad 16/600 Superdex 75 column) in 50mM Tris pH 7.4, 150mM NaCl. Purified SapB was concentrated to 16mg/mL and stored at 4 °C.

His_6_-tagged SapB was expressed in HEK293F cells by polyethylenimine-based transient transfection. After 4 days, SapB-H_6_ was purified from conditioned media by nickel affinity and size exclusion chromatography with a Superdex 75 column (GE Healthcare) in cross-linking buffer. Prior to crystallisation trials (but not cross-linking assays), the tag was removed by incubation with 1.2 U/mL Carboxypeptidase A-agarose at 25°C overnight followed by incubation with Ni-NTA resin to remove tagged protein. The supernatant containing untagged SapB was separated from Carboxypeptidase A-agarose and Ni-NTA agarose by passing through a gravity flow column.

### α-GalCer loading

Endogenously purified lipids were exchanged for the lipid α-GalCer (Avanti Lipids) based on a protocol used to obtain crystal structures of α-GalCer-loaded CD1d^40,41^. Lipid was resuspended in DMSO at 1 mg/mL and dissolved by heating the solution at 80°C for 10 min, with brief sonication and vortexing every 3 minutes. CD1d-β_2_m and SapB were incubated with a 3-fold molar excess of α-GalCer in PBS overnight at room temperature. α-GalCer-loaded proteins were then separated from free α-GalCer by size exclusion chromatography with a Superdex 200 10/300 column equilibrated in 20 mM HEPES pH 7.0, 150 mM NaCl. α-GalCer-loaded proteins were used within 24 hours after size exclusion chromatography to avoid significant hydrolysis of the lipid.

### Equilibrium binding assay

Ni-NTA agarose beads were loaded with a saturating amount of CD1d-β_2_m, and complete capture of CD1d-β_2_m was verified by measuring the absorbance of the supernatant at 280 nm. Saturated beads were washed three times in equilibrium assay buffer (50 mM HEPES pH 7.0, 150 mM NaCl, 0.05% digitonin, 2.5 mM CaCl_2_, 2.5 mM MgCl_2_). Serial dilutions (1:2) of CD1d-β_2_m-saturated beads were made in equilibrium assay buffer supplemented with unloaded Ni-NTA beads to equalize the bead volume across all samples. SapB was added at a final concentration of 680 nM in equilibrium assay buffer. The reaction was incubated with shaking at room temperature for 1 hour. The supernatant containing unbound SapB was separated from the beads by centrifugation (800*g*, 2 min) followed by separation with a Micro-Spin column (Pierce) (800*g*, 2 min) to remove all beads. The supernatant was analysed by SDS-PAGE followed by staining with SYPRO Ruby. For analysis of bead samples, these were boiled in SDS-loading dye, separated by SDS-PAGE and detected by Coomassie staining.

### Cross-linking assay

Amine-reactive cross-linking agents disuccinimidyl sulfoxide (DSSO) and PEGylated bis(sulfosuccinimidyl)suberate (BS(PEG)_5_) were dissolved in DMSO and added to 10 μM CD1d-β_2_m alone, 20 μM SapB alone or 10 μM CD1d + 20 μM SapB with final cross-linker concentrations of 0, 200 μM or 1 mM in cross-linking buffer. The reaction was incubated for 30 min at room temperature and terminated by incubation with 20 mM Tris pH 8 for 15 min. The reaction products were analysed by SDS-PAGE followed by Coomassie staining.

### Crystallography

Crystallization experiments were performed in 96-well nanolitre-scale sitting drops (200 nL protein plus 200 nL of precipitant) equilibrated at 20 °C against 80 μL reservoirs of precipitant. The crystals reported here were grown in drops containing the following components:

Crystal form (i) (Fig. 4) contained glycosylated CD1d-H_6_-β_2_m (13.5 mg/mL) + 2-fold molar excess of unglycosylated SapB (4.7 mg/mL) in 150 mM NaCl, 50 mM Tris pH 7.4 and grew against a reservoir containing 20% w/v Polyethylene glycol 6000, 0.1 M HEPES pH 7 and 0.2 M calcium chloride.

Crystal form (ii) (Fig. 4) contained glycosylated CD1d-H_6_-β_2_m (12.4 mg/mL) + 2-fold molar excess of glycosylated SapB (after tag removal) (4.3 mg/mL) in 20 mM HEPES pH 7.0, 150 mM NaCl and grew against a reservoir containing 20 mM 1,6-Hexanediol, 20 mM 1-Butanol, 20 mM 1,2-Propanediol, 20 mM 2-Propanol, 20 mM 1,4-Butanediol, 20 mM 1,3-Propanediol, 0.1 M Bicine/Trizma base pH 8.5, 12.5% w/v PEG 1000, 12.5% w/v PEG 3350 and 12.5% v/v MPD.

**Table 1.** Data collection statistics and molecular replacement solutions. For Crystal 4B (iii) statistics have been included for data from a single crystal before (isotropic) and after anisotropic truncation and scaling using STARANISO. Values in parentheses are for the highest-resolution shell.

| Data collection | Crystal Fig 4B (iii) |  | Crystal Fig. 4B (iv) | Crystal Fig. 4D (v) |
| --- | --- | --- | --- | --- |
| Beamline | I04 |  | I04-1 | I03 |
| Wavelength (Å) | 0.9999 |  | 0.9159 | 0.9793 |
| Processing package | STARANISO |  | XIA2 3dii | XIA2 DIALS |
| Space group | *C* 2 |  | *P* 2<sub>1</sub> | *C* 2 |
| Cell dimensions |  |  |  |  |
| *a*,*b*,*c* (Å) | 273.3, 88.0, 127.9 |  | 123.8, 122.9, 159.9 | 186.2, 57.1, 129.2 |
| *α*,*β*,*γ* (°) | 90, 114, 90 |  | 90, 95, 90 | 90, 100, 90 |
|  | *Isotropic* | *Anisotropic* |  |  |
| Resolution (Å) | 124.71 – 4.97<br>(5.05 – 4.97) | 124.71 – 4.55<br>(4.99 – 4.55) | 123.27 – 3.49<br>(3.55 – 3.49) | 54.53 – 2.52<br>(2.56 – 2.52) |
| *R*<sub>merge</sub> | 0.438 (2.187) | 0.198 (0.507) | 0.161 (1.572) | 0.072 (2.438) |
| *R*<sub>pim</sub> | 0.279 (1.366) | 0.127 (0.323) | 0.067 (0.663) | 0.031 (1.028) |
| CC<sub>1/2</sub> | 0.967 (0.322) | 0.987 (0.704) | 0.992 (0.714) | 0.999 (0.813) |
| *I*/*σI* | 1.9 (0.5) | 3.4 (1.8) | 6.9 (1.0) | 10.9 (0.8) |
| Completeness (%) | 99.8 (99.8) | e* 80.0 (56.7)<br>s* 39.2 (8.1) | 99.8 (99.9) | 99.8 (98.0) |
| Redundancy | 3.4 (3.5) | 3.4 (3.4) | 6.9 (6.6) | 6.7 (6.6) |
| **Molecular replacement** |  |  |  |  |
| # CD1-β<sub>2</sub>m/ASU | 4 |  | 6 | 2 |

\* ellipsoidal (e) and spherical (s) completeness are quoted for anisotropic data processing using STARANISO.

Crystal form (iii) (Fig. 4) contained α-GalCer-loaded glycosylated CD1d-H_6_-β_2_m (11.9 mg/mL) + 2-fold molar excess of α-GalCer-loaded unglycosylated SapB (4.2 mg/mL) in 50 mM Tris pH 7.4, 150 mM NaCl and grew against a reservoir containing 0.1 M Sodium acetate pH 5.0, 20% w/v PEG 6000 and 0.2 M ammonium chloride.

Crystal form (iv) (Fig. 4) contained deglycosylated CD1d-H_6_-β_2_m (13.3 mg/mL) + 2-fold molar excess of unglycosylated SapB (4.7 mg/mL) in 20 mM citrate pH 6, 150 mM NaCl, and grew against a reservoir containing 16% v/v PEG 500MME, 8% w/v PEG 20000, 0.1 M Bicine/Trizma base pH 8.5, 30 mM Sodium nitrate, 30 mM Sodium phosphate dibasic and 30 mM Ammonium sulfate.

Crystal form (v) (Fig. 4) contained deglycosylated CD1d-H_6_-β_2_m (6.5 mg/mL) + 2-fold molar excess of unglycosylated SapB (2.3 mg/mL) in 100 mM citrate pH 6.0, 150 mM NaCl and grew against a reservoir containing 20% w/v PEG 3350, 0.1 M Bis-Tris propane pH 7.5 and 0.2 M di-Sodium malonate.

### X-ray data collection and molecular replacement

Crystals were cryoprotected in reservoir solution supplemented with 20% (v/v) glycerol and flash-cooled by plunging into liquid nitrogen. Diffraction data were recorded at Diamond Light Source (DLS) on beamlines I03, I04 and I04-1 (Table 1). Diffraction data were indexed and integrated using the automated data processing pipelines at DLS^42^ implementing XIA2 DIALS^43^, XIA2 3dii or STARANISO^44^ depending on the extent of anisotropic diffraction (Table 1), then scaled and merged using AIMLESS^45^. The model for molecular replacement was generated using the published structure of CD1d-β_2_m bound to lysophosphatidylcholine^46^(PDB ID 3U0P, chains A and B), using phenix.sculptor^47,48^ to remove the lipid ligand from the model and revert the glycosylation site mutations present in this structure. Molecular replacement was performed using phenix.phaser^49^ and was followed by rigid body refinement using phenix.refine^50^. Molecular replacement into the highly anisotropic data (Crystal 4Biii) was carried out using both the isotropic and anisotropically processed data resulting in the same solution. Maps were inspected using COOT^51^. Structure figures were rendered using PyMOL (Schrödinger LLC).

### SDS-PAGE analysis of crystals

For each crystallisation condition tested, several crystals were harvested from the crystallisation drop and washed by transferring to a drop of reservoir solution to remove uncrystallised protein components. Crystals were then transferred to a second drop of reservoir solution and dissolved in SDS-PAGE loading buffer. Alternatively, large crystals from which data had been collected at the synchrotron were recovered into SDS-PAGE loading buffer from the crystal-mounting loop. Individual crystal components were analysed by SDS-PAGE followed by Coomassie staining.

## Results

### Purification of CD1d-β_2_m complex from a human cell line

Several strategies have been developed previously for the purification of CD1d including refolding from *E.coli* inclusion bodies^52^, baculovirus-insect cell expression^46^ and mammalian expression systems^53^. An important consideration for CD1d protein expression and purification is the inclusion of the essential partner protein β_2_m. In a manner similar to the heavy chain of MHC class I, CD1d non-covalently associates with β_2_m and this association is required for maturation of its four glycan chains^54^. In order to obtain high quality, near-native CD1d protein for *in vitro* studies, we chose to co-express human CD1d with β_2_m in the mammalian suspension cell line HEK293F as this would confer correct folding, native post-translational modification and full glycosylation of CD1d. Initial trials using polyethylenimine-based transient co-transfection of CD1d and β_2_m resulted in very low yields. Thus we used a piggyBac-expression system^37^ to establish stable and inducible co-expression of full-length, untagged β_2_m with the ectodomain of CD1d possessing a C-terminal hexahistidine tag (H_6_) (Fig. 1A). Both proteins were expressed with an N-terminal secretion signal sequence targeting synthesis to the endoplasmic reticulum resulting in protein secretion into the media. CD1d-H_6_ was purified from conditioned media by nickel affinity chromatography followed by size exclusion chromatography (Fig. 1B and C). Untagged β_2_m co-purified with CD1d, indicating that both proteins are correctly folded and form a stable complex. Although some CD1d alone was purified, we were able to separate this from CD1d-β_2_m following size exclusion (Fig. 1C). Using this expression system, we obtained a high yield (1 mg/100 mL media) of very pure CD1d-β_2_m protein. The yield was over 10 times higher than that obtained with transient co-transfection in the same cell line.

**Figure 1.**
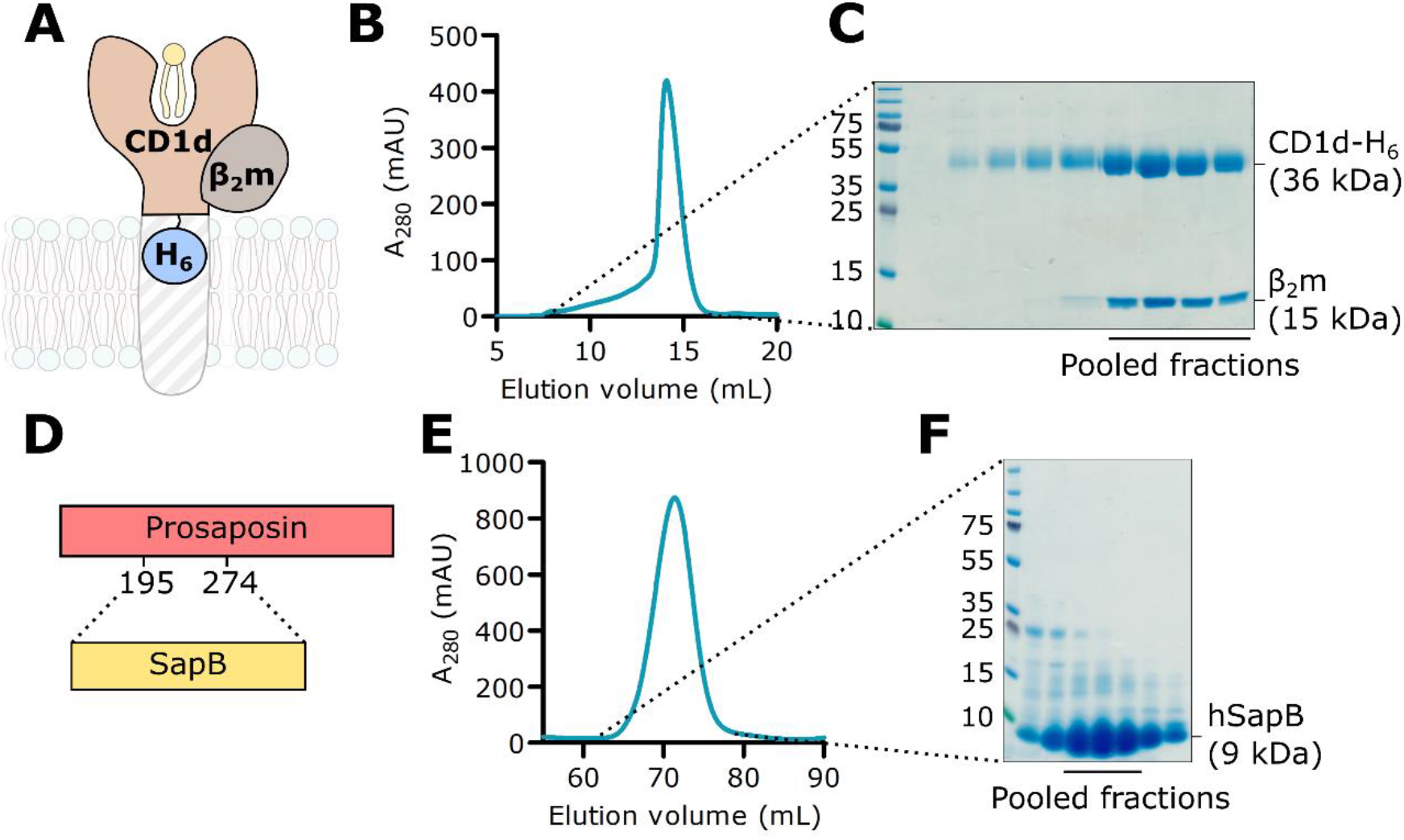
Purification of CD1d-β_2_m and SapB. **A.** A His-tagged construct encoding the extracellular domain of CD1d (CD1d-H_6_) was co-expressed with untagged full-length β_2_m in mammalian cells. **B**. Size exclusion chromatography (SEC) showing co-purification of CD1d-H_6_ with β_2_m. **C**. SDS-PAGE of SEC fractions. The indicated fractions containing β_2_m-associated CD1d were used in subsequent experiments. **D**. Untagged SapB (residues 195-274 of the precursor protein PSAP) was expressed in *E. coli*. **E**. SEC of SapB after multi-step purification. **F**. SDS-PAGE of SEC fractions. The indicated fractions were used in subsequent experiments.

### Purification of SapB

The four saposin proteins (SapA-D) are produced from the proteolytic cleavage of the precursor protein PSAP which normally occurs upon delivery to the lysosome. To produce recombinant SapB alone, the segment of human PSAP corresponding to SapB (Fig. 1D) was expressed in *E. coli*. Untagged SapB was purified following a well-established protocol^39^ that exploits the heat stability of saposins followed by anion exchange and size exclusion chromatography (Fig. 1E and F). Previous structural studies with *E.coli*-expressed SapB identified that it exists as a tight dimer that co-purifies with *E.coli* phospholipids^14^.

### CD1d binding to SapB is not detectable using *in vitro* interaction assays

Classical pull-down experiments involve the capture and immobilisation onto resin of a tagged bait protein followed by incubation with prey protein. Subsequent wash steps remove unbound protein leaving behind protein complexes for detection. However, as the concentration of free prey protein is reduced at each wash, the equilibrium is pushed towards complex dissociation, meaning this method is not well suited for the capture of weak or transient interactions. Using this approach, no interaction was detectable between immobilised CD1d-β_2_m and SapB. Therefore, an alternative approach was adopted to overcome the limitations of prey-depletion by using an *in vitro* equilibrium binding assay (Fig. 2A). In this assay, the bait is bound to beads at a range of concentrations and the prey is added at a low, limiting concentration. Following incubation, and no wash steps, the supernatant is separated from the beads and unbound prey is detected using the sensitive protein dye Sypro Ruby. Monitoring the depletion of prey from the supernatant by the large excess of bait protein, rather than monitoring complex formation directly, represents a more sensitive measure of binding and thus is optimised for weak or transient interactions^55^.

**Figure 2.**
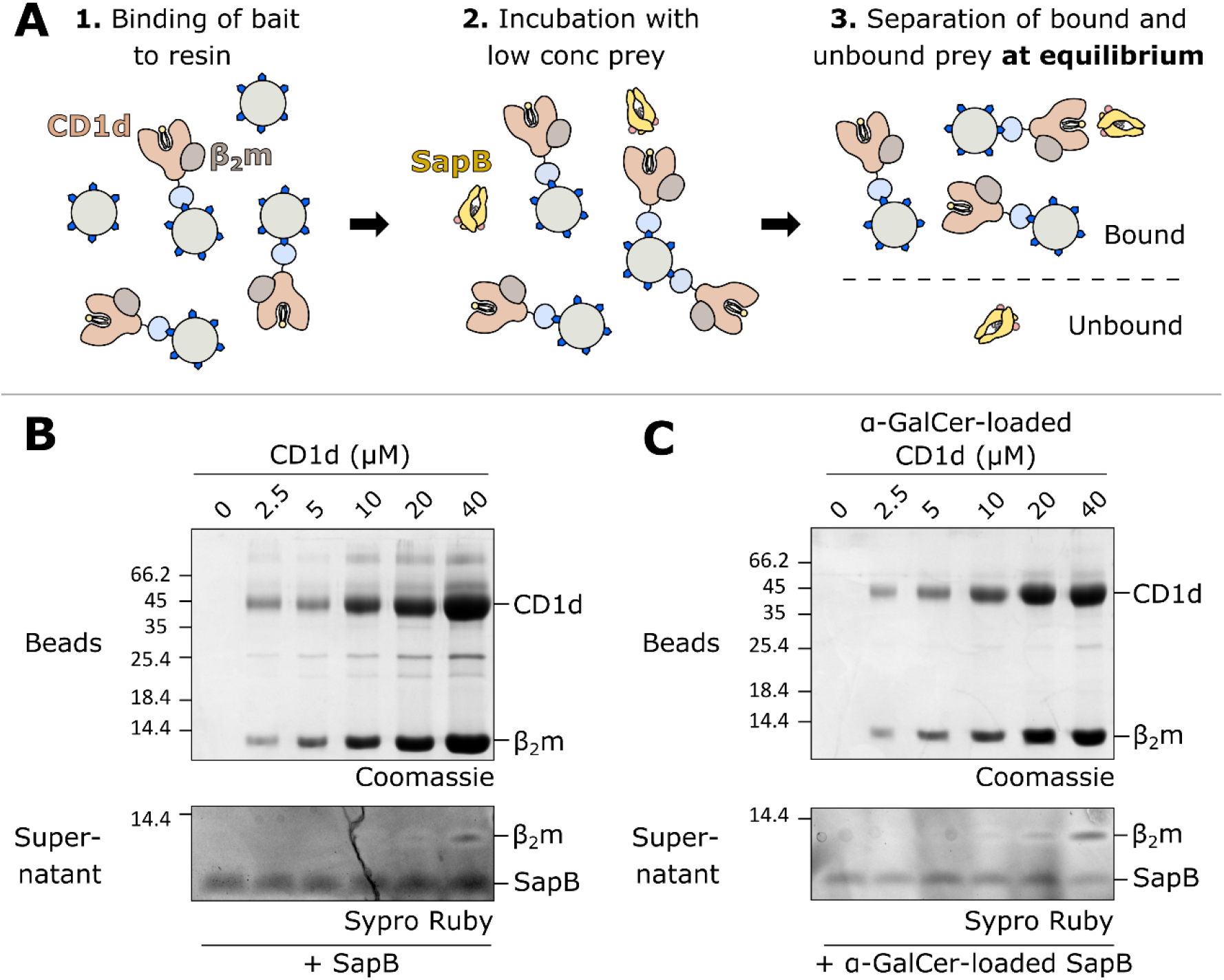
Equilibrium binding assay of CD1d-β_2_m and SapB. **A.** Schematic diagram illustrating the equilibrium binding assay protocol. **B**. Increasing concentrations of CD1d-β_2_m were bound to beads and detected by Coomassie-stained SDS-PAGE. After incubation with SapB, unbound SapB was detected in supernatants by Sypro Ruby-stained SDS-PAGE. **C**. As for panel B but with CD1d-β_2_m and SapB both pre-loaded with the lipid α-GalCer.

Equilibrium binding assays were carried out using CD1d-β_2_m as bait at a concentration range of 0-40 μM and with SapB as the prey at 680 nM. Previous *in vitro* studies have identified that SapB can load α-GalCer onto CD1d at a pH range between 5 and 8^33^. Although this exchange appears to be most efficient at pH 6, this pH causes partial dissociation of the CD1 His-tag from the Ni-NTA resin. Therefore, the equilibrium binding assay was performed at pH 7. Previous work, and our preliminary experiments, showed that β_2_m dissociates from CD1d in the presence of detergent^56^. However, the gentle detergent digitonin was found to cause minimal disruption of the CD1d-β_2_m complex^56^ and was used in this assay. Despite the use of this optimised approach, no binding of SapB to CD1d could be detected (Fig. 2B). This result suggests that CD1d and SapB do not directly interact *in vitro*, or that their interaction is extremely weak with a *K*_d_ > 40 μM. In these conditions, at high CD1d-β_2_m concentrations, a small amount of β_2_m is detectable in the supernatant due to dissociation from CD1d (Fig. 2B). By comparing how little appears to be lost based on the Coomassie-stained bead samples this highlights the sensitivity of this technique and of the Sypro Ruby stain for protein detection.

Previous studies have shown that the ectodomain of CD1d co-purifies with cellular lipids such as sphingomyelin^8–10^, and SapB is known to co-purify with *E. coli* lipids^14^. It is possible that this lipid repertoire could be interfering with direct interactions between these protein components. The lipid α-GalCer has been shown to be loaded onto soluble CD1d by SapB *in vitro*^33,34^, therefore both proteins were pre-loaded with α-GalCer prior to performing the equilibrium binding assay. However, this variation also did not result in a detectable interaction between SapB and CD1d (Fig. 2C).

### *In vitro* cross-linking does not capture an interaction between CD1d and SapB

Cross-linking of proteins using short chemical tethers that react with surface-exposed lysine side-chains can be used to capture transient or weak interactions. Both CD1d and SapB have several surface-exposed lysines, and of particular importance, lysine residues are present near the lipid-binding groove of CD1d where we would predict SapB to bind (Fig. 3A and B). Therefore, it is possible that CD1d and SapB interact in a way that is amenable to intermolecular cross-linking of surface lysines by amine-reactive cross-linkers. We used cross-linkers of different lengths to maximise the chances of capturing an interaction: DSSO, with a 10.3 Å spacer arm, and BS(PEG)_5_, with a 21.7 Å spacer arm. CD1d-β_2_m and SapB were incubated with these cross-linkers either alone or following mixing (Fig. 3C). For the CD1d-β_2_m samples alone, cross-linking between CD1d and β_2_m is clearly evident, as expected. There is also some evidence of additional higher order oligomers, probably due to the large number of surface lysines on both CD1d and β_2_m. However, no additional cross-linking products were observed when CD1d-β_2_m and SapB were incubated together with the cross-linkers. This indicates that CD1d-β_2_m and SapB do not interact together in a conformation that is amenable to cross-linking.

**Figure 3.**
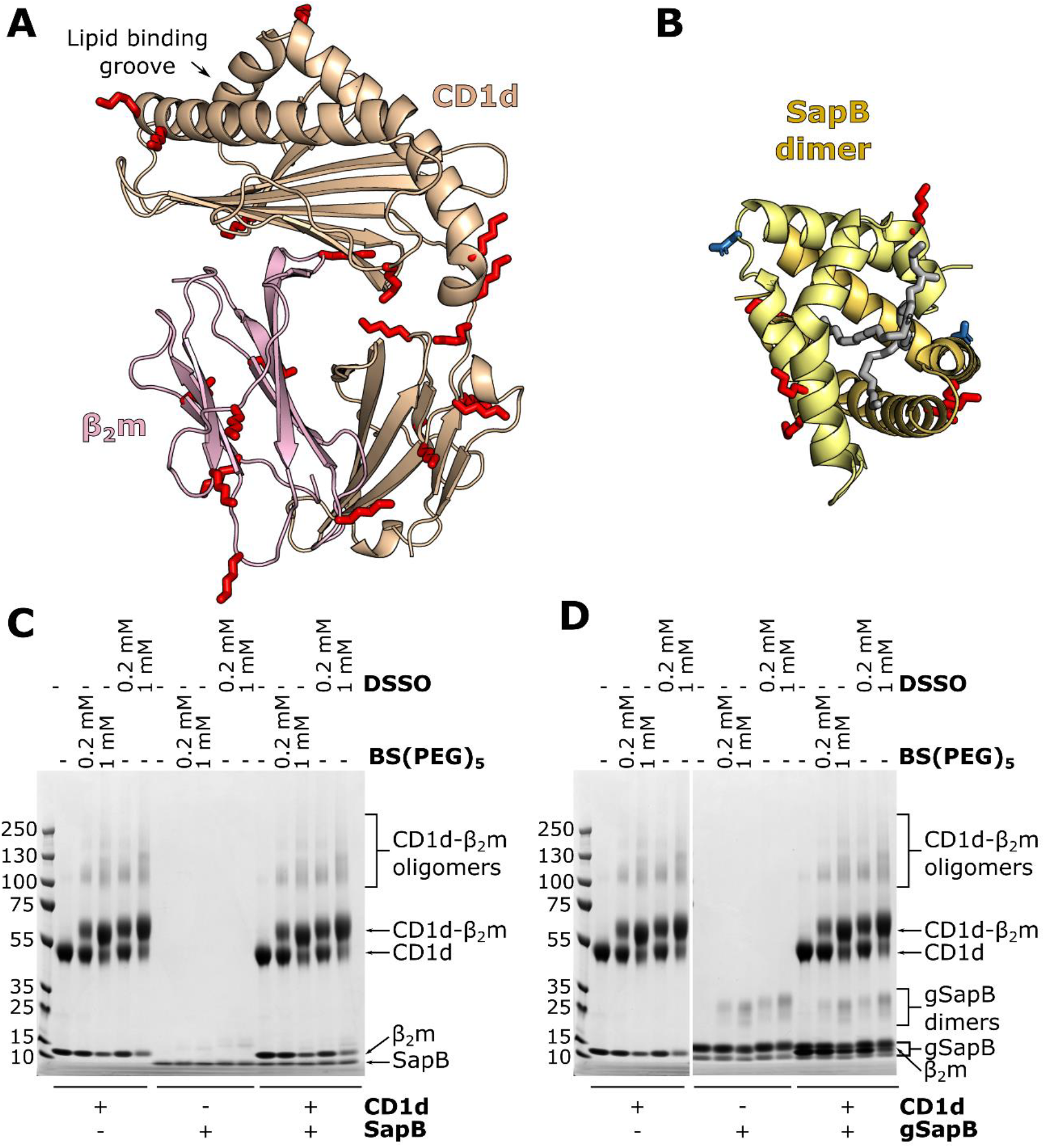
*In vitro* cross-linking of CD1d-β_2_m and SapB. **A.** Ribbon diagram of the structure of CD1d (beige) and β_2_m (pink)(PDB ID 3U0P) with the lipid antigen groove labelled (arrow). Surface-accessible lysine residues available for cross-linking are highlighted (red sticks). **B**. Ribbon diagram of the structure of a SapB dimer (yellow and orange, PDB ID 1N69) with bound lipid (in gray). Lysine residues are highlighted as in panel A. The N-linked glycosylation site of SapB is highlighted (blue sticks). **C**. Coomassie-stained SDS-PAGE of CD1d-β_2_m and unglycosylated SapB samples following incubation with cross-linkers of different lengths: DSSO (10.3 Å) or BS(PEG)_5_ (21.7 Å). **D**. As for panel C but using CD1d-β_2_m and glycosylated SapB (gSapB).

SapB possesses one N-linked glycosylation site on residue Asn21 (Fig. 3B), which remains unglycosylated when expressed in *E.coli*. It is possible that this glycan on SapB may contribute to its interaction with CD1d. Therefore, we repeated the cross-linking experiment following expression of His-tagged SapB in mammalian HEK293F cells, enabling the purification of glycosylated protein. Following purification, two SapB-H_6_ species were present, with molecular weights of approximately 10 and 13 kDa, likely corresponding to unglycosylated and glycosylated SapB-H_6_ (Fig. 3D). It was not possible to separate these two SapB species, likely due to SapB dimerization, and so cross-linking experiments were carried out using this mixed sample (Fig. 3D). However, we again did not observe formation of any additional cross-linked products when CD1d-β_2_m was incubated with glycosylated SapB-H_6_. We note that although the same amount of unglycosylated SapB (Fig. 3C) and glycosylated SapB (Fig. 3D) were used in cross-linking assays, the Coomassie-stained bands corresponding to glycosylated SapB appear much stronger. This increased staining may be due to the additional histidine residues in the affinity tag increasing the amount of Coomassie reagent binding to this small protein.

### CD1d-β_2_m and SapB do not co-crystallise

An alternative method to capture weak, transient protein-protein interactions is to attempt co-crystallisation. In this experiment both protein components are mixed together at a range of concentrations and different molar ratios, then subjected to crystallisation trials. The benefit of using the sitting-drop vapour diffusion approach is that during the experiment the drop containing protein and precipitant gradually equilibrates with the reservoir, generally resulting in reduced drop volume and the progressive increase in concentration of the drop components. In this way, the protein components are steadily concentrated together with the aim of capturing an interaction that is otherwise extremely hard to measure using biochemical and biophysical assays^57^.

Extensive crystallisation trials containing both CD1d-β_2_m and SapB were performed, varying the following parameters: (1) protein concentration (6-13 mg/mL CD1d-β_2_m, 2-6 mg/mL SapB), (2) molar ratio of SapB to CD1d, taking into account the predicted dimeric state of SapB (2:1-3:1 SapB to CD1d); (3) full glycosylation or deglycosylation of protein components; and (4) lipid exchange for α-GalCer. Figure 4A shows an example of the high purity of the protein components that went into crystallisation trials. These experiments yielded several different crystals in a range of conditions, a subset of which are shown in Figure 4B. To determine the composition of these crystals, two approaches were used depending on the quality of the crystals grown. The first approach, which can be used for any protein crystal, involves harvesting and washing the crystals in reservoir solution to remove residual uncrystallised components, then separating the proteins by SDS-PAGE followed by detection using Coomassie staining (Fig. 4C, each gel lane corresponds to the above crystal in panel 4B). This revealed that both SapB and CD1d-β_2_m were capable of forming crystals but in none of the crystals grown did both components crystallise together. The second approach used was to collect diffraction data for any crystals that diffracted to 5 Å resolution or better, then use molecular replacement to confirm the contents of the asymmetric unit (Table 1). Three crystal forms satisfied this criteria, including two that had also been tested using SDS-PAGE (Fig. 4B (iii) and (iv) and 4D (v)). These three samples all contained CD1d-β_2_m only as confirmed by molecular replacement trials using multiple copies of all potential components as well as manual inspection of maps. In all datasets additional density was observed for unmodelled glycans on CD1d (deglycosylation of CD1d using Endo H leaves a residual acetylated glycosamine), confirming the validity of the MR solution. For several CD1d molecules, additional unmodelled density near the lipid-binding groove was evident that likely corresponds to partially-ordered lipid components. Further inspection of electron density maps revealed no evidence for the presence of SapB. For most molecules of CD1d in all three different crystal forms the predicted binding site for SapB (over the lipid-binding groove) was occupied by another copy of CD1d. The use of both SDS-PAGE analysis of all grown crystals and MR solution of higher quality crystals confirmed that, despite extensive efforts with a broad range of CD1d-β_2_m/SapB combinations, these do not form a stable, crystallisable complex.

**Figure 4.**
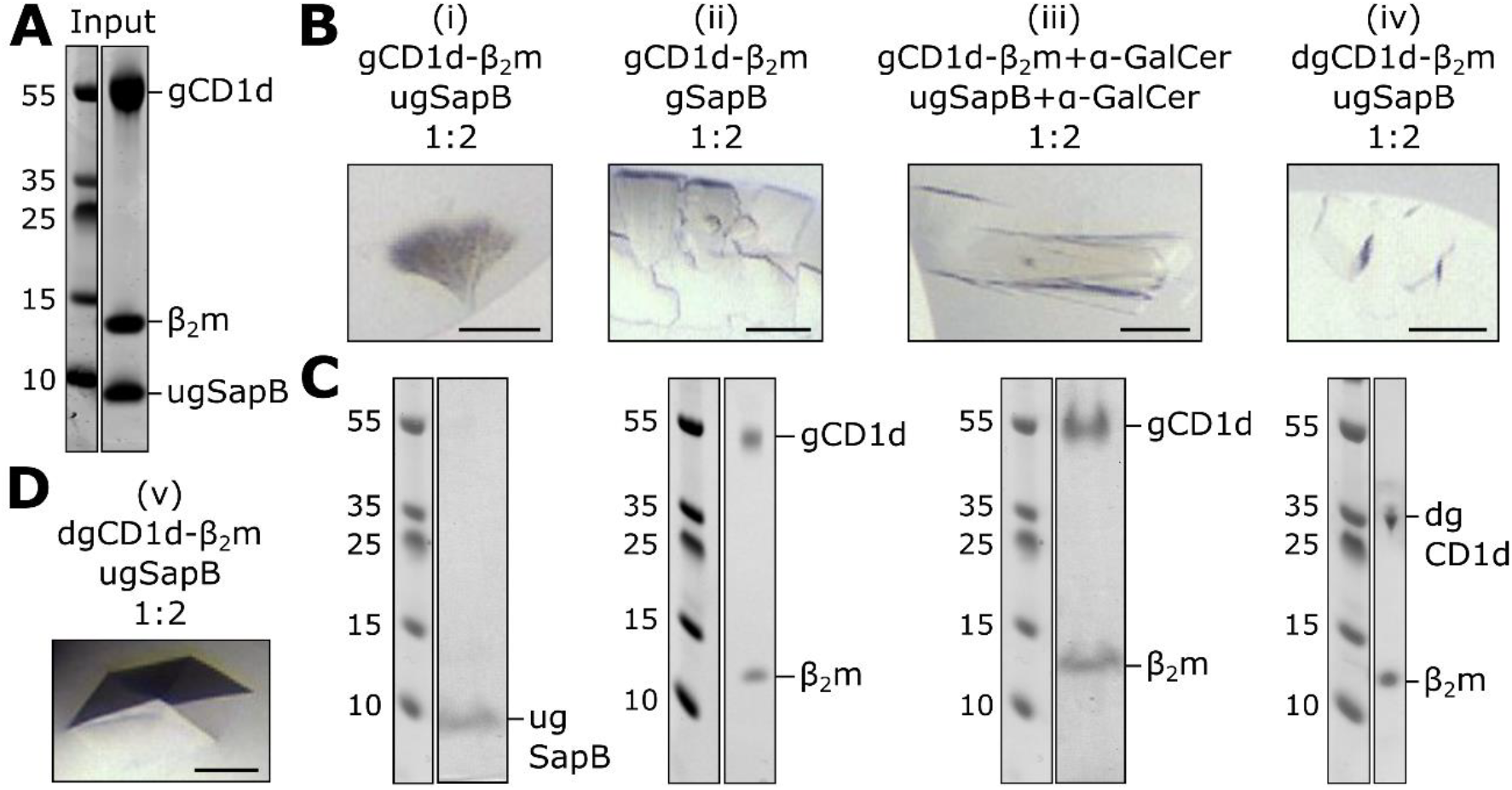
Crystallisation trials of CD1d-β_2_m and SapB. **A.** Coomassie-stained SDS-PAGE example of the purified protein sample used in crystallisation trials. **B.** Examples of four crystals (i)-(iv) grown in a range of conditions using different protein compositions. Scale bar represents 100 μm. **C.** Coomassie-stained SDS-PAGE of harvested and washed crystals shown in panel B. **D.** Image of the crystal that diffracted to highest resolution as described in Table 1. Scale bar represents 100 μm. Abbreviations: gCD1d-glycosylated CD1d, dgCD1d-deglycosylated CD1d, gSapB-glycosylated SapB, and ugSapB-unglycosylated SapB.

## Discussion

Using highly purified components we have been unable to identify a direct interaction between CD1d-β_2_m and SapB *in vitro*. Several biochemical and structural approaches were implemented that have the capacity to capture weak and transient interactions, including binding equilibrium assays, protein cross-linking, and co-crystallisation trials. Various factors that may influence the strength or specificity of these interactions were tested, including different pH and buffer systems, exchange of the lipids loaded onto both protein components, and changing the glycosylation state of each protein. None of these variants resulted in the detection of a direct interaction. Our previous work studying saposin interactions with lysosomal hydrolases showed that although these interactions are sensitive to pH and salt concentrations, they were always detectable, and while detergent/lipid was required, neither the nature of the lipid/detergent nor the glycosylation state mattered^19^.

The study of lipid loading of CD1 molecules is technically challenging as biochemical methods often require the inclusion of detergents, which may interfere with lipid solubilisation and binding. Also, the detection of changes to CD1d-lipid complexes is limited by available reagents including specific antibodies and iNKT TCRs. However, the inability to identify a direct interaction between CD1d and SapB raises a number of important questions regarding the role of saposins in lipid antigen presentation.

It is well established that knockout of the saposin precursor protein PSAP results in impaired presentation of lipid antigens by CD1d. Due to the dual role of saposins in lipid loading and lipid processing, the absence of PSAP will not only alter CD1d lipid loading, but will also significantly alter the composition of the lysosomal lipidome. Indeed, the loss of a single saposin results in the cellular accumulation of multiple, different GSLs. For example, loss of SapB alone, in mice or in humans, results in the accumulation of sulfatides, globotriaosylceramide, and lactosylceramide^58,59^. Therefore, in saposin knockout and rescue models, the effects on antigen processing and the changes to local lipid concentrations may have indirect and unpredictable effects on CD1d loading in the lysosome. Similarly, cell-based assays that involve recognition of CD1d-lipid complexes by NKT cells are made complicated by the requirement of some glycolipid headgroups to be processed by lysosomal hydrolases^12^, a process that is itself dependent upon saposins.

*In vitro* assays testing the role of different saposins in facilitating lysosomal hydrolase activity identify some of the difficulties in determining specificity of saposin-lipid binding. For example, *in vitro* assays have identified that SapC and SapA are both able to enhance the activity of β-galactosylceramidase (GALC) to promote degradation of the galactosphingolipids^60,61^. However, this does not reflect the specificity of saposins in whole organisms as loss of SapA causes Krabbe disease, similar to that of loss of GALC, while mutations in SapC do not^20,21,23–25^. Why these *in vitro* assays do not recapitulate the specificity of saposins in a whole organism remains unclear but may suggest a non-specific detergent-like effect of saposins when present at relatively high concentrations. It is unclear whether a similar effect may be partially confounding the interpretation of SapB influence on CD1d lipid loading. While *in vitro* studies demonstrating the ability of SapB to enhance CD1d stimulation of iNKT cells or binding to iNKT TCRs are very compelling regarding the importance of SapB in this process, these do represent an indirect way of monitoring how SapB may be involved in lipid exchange^33,34^. In light of our data showing no detectable interaction between CD1d and SapB, the question remains: how can SapB facilitate lipid editing without directly binding CD1d? Modelling of how SapB might facilitate lipid exchange identified a role for SapB in enhancing the kinetics of lipid exchange in a manner similar to that of Tapasin in peptide exchange on MHC class-I molecules^34^. In order for Tapasin to carry out this role, it does bind directly to MHC class-I but as part of a larger complex including additional proteins^2,62^. Although the *in vitro* assays demonstrating the importance of SapB for CD1d lipid presentation did not require other proteins, it is possible that there may be additional factors involved in enhancing the affinity of SapB for CD1d in cells.

Additional factors may influence the efficiency of lipid loading onto CD1d in cells. For example, the relative abundance of different LTPs in the lysosome may influence the lysosomal lipid composition, altering exchange kinetics. Therefore, it may be important that SapA-D exist in equimolar concentrations in the endolysosome, as ensured by their production from a common precursor. If this is the case, studies involving individual saposins or partial PSAP rescues will be difficult to interpret, potentially explaining some experimental inconsistencies. The lysosomal lipidome is not only influenced by the activity of saposins and sphingolipid processing hydrolases but also by the careful balance of other membrane components such as cholesterol and bis(monoacylglycero)phosphate that are crucial for endosomal vesicle formation. Lipid loading of CD1 in endolysosomal compartments is highly complex, requiring the delicate interplay of lipid binding, lipid processing and lipid exchange. For now, the mechanism of transfer of lipids onto CD1 molecules, as well as the molecular determinants of LTP binding specificity, remains unclear.

### Grant information

M.S. is supported by a Wellcome Trust PhD studentship. S.C.G. is supported by a Sir Henry Dale Fellowship co-funded by the Royal Society and Wellcome Trust (098406/Z/12/B). J.E.D. is supported by a Royal Society University Research Fellowship (UF100371).

## Acknowledgements

We acknowledge Diamond Light Source for time on beamlines I03, I04 and I04-1 under proposal MX15916. Remote access was supported in part by the EU FP7 infrastructure grant BIOSTRUCT-X (Contract No. 283570).

## References

1. Pos, W., Sethi, D.K., Call, M.J., Schulze, M.S., Anders, A.K., Pyrdol, J. & Wucherpfennig, K.W. Crystal structure of the HLA-DM-HLA-DR1 complex defines mechanisms for rapid peptide selection. Cell 151, 1557–68 (2012).

2. Blees, A., Januliene, D., Hofmann, T., Koller, N., Schmidt, C., Trowitzsch, S., Moeller, A. & Tampe, R. Structure of the human MHC-I peptide-loading complex. Nature 551, 525–528 (2017).

3. Ilca, F.T., Neerincx, A., Hermann, C., Marcu, A., Stevanovic, S., Deane, J.E. & Boyle, L.H. TAPBPR mediates peptide dissociation from MHC class I using a leucine lever. Elife 7(2018).

4. Layre, E., de Jong, A. & Moody, D.B. Human T cells use CD1 and MR1 to recognize lipids and small molecules. Curr Opin Chem Biol 23, 31–8 (2014).

5. Moody, D.B. & Cotton, R.N. Four pathways of CD1 antigen presentation to T cells. Curr Opin Immunol 46, 127–133 (2017).

6. Brennan, P.J., Brigl, M. & Brenner, M.B. Invariant natural killer T cells: an innate activation scheme linked to diverse effector functions. Nat Rev Immunol 13, 101–17 (2013).

7. Sprong, H., van der Sluijs, P. & van Meer, G. How proteins move lipids and lipids move proteins. Nat Rev Mol Cell Biol 2, 504–13 (2001).

8. Cox, D., Fox, L., Tian, R., Bardet, W., Skaley, M., Mojsilovic, D., Gumperz, J. & Hildebrand, W. Determination of cellular lipids bound to human CD1d molecules. PLoS One 4, e5325 (2009).

9. Yuan, W., Kang, S.J., Evans, J.E. & Cresswell, P. Natural lipid ligands associated with human CD1d targeted to different subcellular compartments. J Immunol 182, 4784–91 (2009).

10. Haig, N.A., Guan, Z.Q., Li, D.M., McMichael, A., Raetz, C.R.H. & Xu, X.N. Identification of Self-lipids Presented by CD1c and CD1d Proteins. J Biol Chem 286, 37692–37701 (2011).

11. Teyton, L. Role of lipid transfer proteins in loading CD1 antigen-presenting molecules. J Lipid Res 59, 1367–1373 (2018).

12. Prigozy, T.I., Naidenko, O., Qasba, P., Elewaut, D., Brossay, L., Khurana, A., Natori, T., Koezuka, Y., Kulkarni, A. & Kronenberg, M. Glycolipid antigen processing for presentation by CD1d molecules. Science 291, 664–7 (2001).

13. Darmoise, A., Teneberg, S., Bouzonville, L., Brady, R.O., Beck, M., Kaufmann, S.H. & Winau, F. Lysosomal alpha-galactosidase controls the generation of self lipid antigens for natural killer T cells. Immunity 33, 216–28 (2010).

14. Ahn, V.E., Faull, K.F., Whitelegge, J.P., Fluharty, A.L. & Prive, G.G. Crystal structure of saposin B reveals a dimeric shell for lipid binding. Proc Natl Acad Sci U S A 100, 38–43 (2003).

15. de Alba, E., Weiler, S. & Tjandra, N. Solution structure of human saposin C: pH-dependent interaction with phospholipid vesicles. Biochemistry 42, 14729–40 (2003).

16. Ahn, V.E., Leyko, P., Alattia, J.R., Chen, L. & Prive, G.G. Crystal structures of saposins A and C. Protein Sci 15, 1849–57 (2006).

17. Popovic, K. & Prive, G.G. Structures of the human ceramide activator protein saposin D. Acta Crystallogr D Biol Crystallogr 64, 589–94 (2008).

18. Popovic, K., Holyoake, J., Pomes, R. & Prive, G.G. Structure of saposin A lipoprotein discs. Proc Natl Acad Sci U S A 109, 2908–12 (2012).

19. Hill, C.H., Cook, G.M., Spratley, S.J., Fawke, S., Graham, S.C. & Deane, J.E. The mechanism of glycosphingolipid degradation revealed by a GALC-SapA complex structure. Nat Commun 9, 151 (2018).

20. Matsuda, J., Vanier, M.T., Saito, Y., Tohyama, J., Suzuki, K. & Suzuki, K. A mutation in the saposin A domain of the sphingolipid activator protein (prosaposin) gene results in a late-onset, chronic form of globoid cell leukodystrophy in the mouse. Hum Mol Genet 10, 1191–9 (2001).

21. Spiegel, R., Bach, G., Sury, V., Mengistu, G., Meidan, B., Shalev, S., Shneor, Y., Mandel, H. & Zeigler, M. A mutation in the saposin A coding region of the prosaposin gene in an infant presenting as Krabbe disease: first report of saposin A deficiency in humans. Mol Genet Metab 84, 160–6 (2005).

22. Kuchar, L., Ledvinova, J., Hrebicek, M., Myskova, H., Dvorakova, L., Berna, L., Chrastina, P., Asfaw, B., Elleder, M., Petermoller, M., Mayrhofer, H., Staudt, M., Krageloh-Mann, I., Paton, B.C. & Harzer, K. Prosaposin deficiency and saposin B deficiency (activator-deficient metachromatic leukodystrophy): report on two patients detected by analysis of urinary sphingolipids and carrying novel PSAP gene mutations. Am J Med Genet A 149A, 613–21 (2009).

23. Vaccaro, A.M., Motta, M., Tatti, M., Scarpa, S., Masuelli, L., Bhat, M., Vanier, M.T., Tylki-Szymanska, A. & Salvioli, R. Saposin C mutations in Gaucher disease patients resulting in lysosomal lipid accumulation, saposin C deficiency, but normal prosaposin processing and sorting. Hum Mol Genet 19, 2987–97 (2010).

24. Motta, M., Camerini, S., Tatti, M., Casella, M., Torreri, P., Crescenzi, M., Tartaglia, M. & Salvioli, R. Gaucher disease due to saposin C deficiency is an inherited lysosomal disease caused by rapidly degraded mutant proteins. Hum Mol Genet 23, 5814–26 (2014).

25. Kang, L., Zhan, X., Ye, J., Han, L., Qiu, W., Gu, X. & Zhang, H. A rare form of Gaucher disease resulting from saposin C deficiency. Blood Cells Mol Dis 68, 60–65 (2018).

26. Locatelli-Hoops, S., Remmel, N., Klingenstein, R., Breiden, B., Rossocha, M., Schoeniger, M., Koenigs, C., Saenger, W. & Sandhoff, K. Saposin A mobilizes lipids from low cholesterol and high bis(monoacylglycerol)phosphate-containing membranes: patient variant Saposin A lacks lipid extraction capacity. J Biol Chem 281, 32451–60 (2006).

27. Remmel, N., Locatelli-Hoops, S., Breiden, B., Schwarzmann, G. & Sandhoff, K. Saposin B mobilizes lipids from cholesterol-poor and bis(monoacylglycero)phosphate-rich membranes at acidic pH - Unglycosylated patient variant saposin B lacks lipid-extraction capacity. FEBS J 274, 3405–3420 (2007).

28. Alattia, J.R., Shaw, J.E., Yip, C.M. & Prive, G.G. Direct visualization of saposin remodelling of lipid bilayers. J Mol Biol 362, 943–53 (2006).

29. Alattia, J.R., Shaw, J.E., Yip, C.M. & Prive, G.G. Molecular imaging of membrane interfaces reveals mode of beta-glucosidase activation by saposin C. Proc Natl Acad Sci U S A 104, 17394–9 (2007).

30. Leon, L., Tatituri, R.V., Grenha, R., Sun, Y., Barral, D.C., Minnaard, A.J., Bhowruth, V., Veerapen, N., Besra, G.S., Kasmar, A., Peng, W., Moody, D.B., Grabowski, G.A. & Brenner, M.B. Saposins utilize two strategies for lipid transfer and CD1 antigen presentation. Proc Natl Acad Sci U S A 109, 4357–64 (2012).

31. Zhou, D., Cantu, C., 3rd, Sagiv, Y., Schrantz, N., Kulkarni, A.B., Qi, X., Mahuran, D.J., Morales, C.R., Grabowski, G.A., Benlagha, K., Savage, P., Bendelac, A. & Teyton, L. Editing of CD1d-bound lipid antigens by endosomal lipid transfer proteins. Science 303, 523–7 (2004).

32. Kang, S.J. & Cresswell, P. Saposins facilitate CD1d-restricted presentation of an exogenous lipid antigen to T cells. Nat Immunol 5, 175–81 (2004).

33. Yuan, W., Qi, X., Tsang, P., Kang, S.J., Illarionov, P.A., Besra, G.S., Gumperz, J. & Cresswell, P. Saposin B is the dominant saposin that facilitates lipid binding to human CD1d molecules. Proc Natl Acad Sci U S A 104, 5551–6 (2007).

34. Salio, M., Ghadbane, H., Dushek, O., Shepherd, D., Cypen, J., Gileadi, U., Aichinger, M.C., Napolitani, G., Qi, X., van der Merwe, P.A., Wojno, J., Veerapen, N., Cox, L.R., Besra, G.S., Yuan, W., Cresswell, P. & Cerundolo, V. Saposins modulate human invariant Natural Killer T cells self-reactivity and facilitate lipid exchange with CD1d molecules during antigen presentation. Proc Natl Acad Sci U S A 110, E4753–61 (2013).

35. Schrantz, N., Sagiv, Y., Liu, Y., Savage, P.B., Bendelac, A. & Teyton, L. The Niemann-Pick type C2 protein loads isoglobotrihexosylceramide onto CD1d molecules and contributes to the thymic selection of NKT cells. J Exp Med 204, 841–52 (2007).

36. Winau, F., Schwierzeck, V., Hurwitz, R., Remmel, N., Sieling, P.A., Modlin, R.L., Porcelli, S.A., Brinkmann, V., Sugita, M., Sandhoff, K., Kaufmann, S.H. & Schaible, U.E. Saposin C is required for lipid presentation by human CD1b. Nat Immunol 5, 169–74 (2004).

37. Li, Z., Michael, I.P., Zhou, D., Nagy, A. & Rini, J.M. Simple piggyBac transposon-based mammalian cell expression system for inducible protein production. Proc Natl Acad Sci U S A 110, 5004–9 (2013).

38. Aricescu, A.R., Lu, W. & Jones, E.Y. A time- and cost-efficient system for high-level protein production in mammalian cells. Acta Crystallogr D Biol Crystallogr 62, 1243–50 (2006).

39. Hill, C.H., Read, R.J. & Deane, J.E. Structure of human saposin A at lysosomal pH. Acta Crystallogr F Struct Biol Commun 71, 895–900 (2015).

40. Pellicci, D.G., Patel, O., Kjer-Nielsen, L., Pang, S.S., Sullivan, L.C., Kyparissoudis, K., Brooks, A.G., Reid, H.H., Gras, S., Lucet, I.S., Koh, R., Smyth, M.J., Mallevaey, T., Matsuda, J.L., Gapin, L., McCluskey, J., Godfrey, D.I. & Rossjohn, J. Differential recognition of CD1d-alpha-galactosyl ceramide by the V beta 8.2 and V beta 7 semi-invariant NKT T cell receptors. Immunity 31, 47–59 (2009).

41. Lopez-Sagaseta, J., Kung, J.E., Savage, P.B., Gumperz, J. & Adams, E.J. The molecular basis for recognition of CD1d/alpha-galactosylceramide by a human non-Valpha24 T cell receptor. PLoS Biol 10, e1001412 (2012).

42. Winter, G. xia2: an expert system for macromolecular crystallography data reduction. J Appl Crystallogr 43, 186–190 (2010).

43. Winter, G., Waterman, D.G., Parkhurst, J.M., Brewster, A.S., Gildea, R.J., Gerstel, M., Fuentes-Montero, L., Vollmar, M., Michels-Clark, T., Young, I.D., Sauter, N.K. & Evans, G. DIALS: implementation and evaluation of a new integration package. Acta Crystallogr D Struct Biol 74, 85–97 (2018).

44. Tickle, I.J., Flensburg, C., Keller, P., Paciorek, W., Sharff, A., & Vonrhein, C., Bricogne, G. STARANISO. Cambridge, United Kingdom: Global Phasing Ltd. (2018).

45. Evans, P.R. & Murshudov, G.N. How good are my data and what is the resolution. Acta Crystallogr D Biol Crystallogr 69, 1204–1214 (2013).

46. Lopez-Sagaseta, J., Sibener, L.V., Kung, J.E., Gumperz, J. & Adams, E.J. Lysophospholipid presentation by CD1d and recognition by a human Natural Killer T-cell receptor. EMBO J 31, 2047–59 (2012).

47. Adams, P.D., Afonine, P.V., Bunkoczi, G., Chen, V.B., Davis, I.W., Echols, N., Headd, J.J., Hung, L.W., Kapral, G.J., Grosse-Kunstleve, R.W., McCoy, A.J., Moriarty, N.W., Oeffner, R., Read, R.J., Richardson, D.C., Richardson, J.S., Terwilliger, T.C. & Zwart, P.H. PHENIX: a comprehensive Python-based system for macromolecular structure solution. Acta Crystallogr D Biol Crystallogr 66, 213–21 (2010).

48. Bunkoczi, G. & Read, R.J. Improvement of molecular-replacement models with Sculptor. Acta Crystallogr D Biol Crystallogr 67, 303–12 (2011).

49. McCoy, A.J., Grosse-Kunstleve, R.W., Adams, P.D., Winn, M.D., Storoni, L.C. & Read, R.J. Phaser crystallographic software. J Appl Crystallogr 40, 658–674 (2007).

50. Afonine, P.V., Grosse-Kunstleve, R.W., Echols, N., Headd, J.J., Moriarty, N.W., Mustyakimov, M., Terwilliger, T.C., Urzhumtsev, A., Zwart, P.H. & Adams, P.D. Towards automated crystallographic structure refinement with phenix.refine. Acta Crystallogr D Biol Crystallogr 68, 352–67 (2012).

51. Emsley, P., Lohkamp, B., Scott, W.G. & Cowtan, K. Features and development of Coot. Acta Crystallogr D Biol Crystallogr 66, 486–501 (2010).

52. Koch, M., Stronge, V.S., Shepherd, D., Gadola, S.D., Mathew, B., Ritter, G., Fersht, A.R., Besra, G.S., Schmidt, R.R., Jones, E.Y. & Cerundolo, V. The crystal structure of human CD1d with and without alpha-galactosylceramide. Nat Immunol 6, 819–826 (2005).

53. Gumperz, J.E., Roy, C., Makowska, A., Lum, D., Sugita, M., Podrebarac, T., Koezuka, Y., Porcelli, S.A., Cardell, S., Brenner, M.B. & Behar, S.M. Murine CD1d-restricted T cell recognition of cellular lipids. Immunity 12, 211–21 (2000).

54. Kim, H.S., Garcia, J., Exley, M., Johnson, K.W., Balk, S.P. & Blumberg, R.S. Biochemical characterization of CD1d expression in the absence of beta2-microglobulin. J Biol Chem 274, 9289–95 (1999).

55. Pollard, T.D. A guide to simple and informative binding assays. Mol Biol Cell 21, 4061–7 (2010).

56. Paduraru, C., Spiridon, L., Yuan, W., Bricard, G., Valencia, X., Porcelli, S.A., Illarionov, P.A., Besra, G.S., Petrescu, S.M., Petrescu, A.J. & Cresswell, P. An N-linked glycan modulates the interaction between the CD1d heavy chain and beta 2-microglobulin. J Biol Chem 281, 40369–78 (2006).

57. Muenzner, J., Traub, L.M., Kelly, B.T. & Graham, S.C. Cellular and viral peptides bind multiple sites on the N-terminal domain of clathrin. Traffic 18, 44–57 (2017).

58. Li, S.C., Kihara, H., Serizawa, S., Li, Y.T., Fluharty, A.L., Mayes, J.S. & Shapiro, L.J. Activator protein required for the enzymatic hydrolysis of cerebroside sulfate. Deficiency in urine of patients affected with cerebroside sulfatase activator deficiency and identity with activators for the enzymatic hydrolysis of GM1 ganglioside and globotriaosylceramide. J Biol Chem 260, 1867–71 (1985).

59. Sun, Y., Witte, D.P., Ran, H., Zamzow, M., Barnes, S., Cheng, H., Han, X., Williams, M.T., Skelton, M.R., Vorhees, C.V. & Grabowski, G.A. Neurological deficits and glycosphingolipid accumulation in saposin B deficient mice. Hum Mol Genet 17, 2345–56 (2008).

60. Harzer, K., Paton, B.C., Christomanou, H., Chatelut, M., Levade, T., Hiraiwa, M. & O’Brien, J.S. Saposins (sap) A and C activate the degradation of galactosylceramide in living cells. FEBS Lett 417, 270–4 (1997).

61. Harzer, K., Hiraiwa, M. & Paton, B.C. Saposins (sap) A and C activate the degradation of galactosylsphingosine. FEBS Lett 508, 107–10 (2001).

62. Dong, G., Wearsch, P.A., Peaper, D.R., Cresswell, P. & Reinisch, K.M. Insights into MHC class I peptide loading from the structure of the tapasin-ERp57 thiol oxidoreductase heterodimer. Immunity 30, 21–32 (2009).

